# Burrowing crabs and physical factors hasten marsh recovery at panne edges

**DOI:** 10.1101/2021.03.17.435836

**Authors:** Kathryn M. Beheshti, Charlie Endris, Peter Goodwin, Annabelle Pavlak, Kerstin Wasson

**Affiliations:** Department of Ecology and Evolutionary Biology, University of California, Santa Cruz; Elkhorn Slough National Estuarine Research Reserve; Center for Environmental Science, University of Maryland Center for Environmental Science

**Keywords:** marsh loss, panne dynamics, crabs, plant-animal interactions, central California, *Pachygrapsus crassipes*, *Salicornia pacifica*

## Abstract

Salt marsh loss is projected to increase as sea-level rise accelerates with global climate change. Salt marsh loss occurs along both lateral creek and channel edges and in the marsh interior, when pannes expand and coalesce. Often, edge loss is attributed to erosive processes whereas dieback in the marsh interior is linked to excessive inundation or deposition of wrack. We conducted a two-year field experiment (2016-2018) in a central California estuary, where, immediately preceding our study, marsh dieback at creek edges and panne expansion occurred during a period of severe drought and an overlapping warm water event. Our study explored how an abundant burrowing crab, shown to have strong negative effects on marsh biomass near creek edges, affects panne dynamics. We also explored which panne attributes best predicted their dynamics. Overall, we found that pannes contracted during the study period, but with variable rates of marsh recovery across pannes. Our model incorporating both physical and biological factors explained 86% of the variation in panne contraction. The model revealed a positive effect of crab activity, sediment accretion, and a composite of depth and elevation on panne contraction, and a negative effect of panne size and distance to nearest panne. The positive crab effects detected in pannes contrast with negative effects we had earlier detected near creek edges, highlighting the context-dependence of top-down and bioturbation effects in marshes. As global change continues and the magnitude and frequency of disturbances increases, understanding the dynamics of marsh loss in the marsh interior as well as creek banks will be critical for the management of these coastal habitats.

## INTRODUCTION

Salt marshes are dynamic systems and generally resilient to perturbations, yet their ability to respond to the multitude of stressors they face has been compromised [1,2]. This is largely due to the many human alterations (diversion of freshwater, depleted sediment supply, reclamation, pollution, eutrophication, barriers to marsh migration) that salt marshes have endured over the last two centuries [3]. The degradation of salt marsh habitat is of great concern, especially in the face of accelerating sea-level rise [1,4]. Since the 1800s, it is estimated that 25-90% of salt marsh habitats have been lost [5,6]. Further loss of salt marsh habitat would come at a great cost as these systems are some of the most productive coastal habitats in the world and they support many high-valued ecosystem services (carbon sink, storm buffer, nursery habitat) [7,8].

Loss of vegetation can occur along tidal channel or creek bank edges (hereafter creek edges) and in the marsh interior [9]. Along creek edges, wave erosion can undercut the marsh scarp and lead to erosional events where large sections of marsh are lost [2,10]. Insufficient sediment supply [11,12], low belowground marsh biomass [13], and algal wrack deposition [14] further hasten creek edge retreat. Features that reduce wave fetch and intensity of wave or boat wake action such as oyster reefs can slow salt marsh erosion [15]. Similarly, benthic diatoms secrete extracellular polymeric substances that both enhance sediment accretion and cohesion and reduce erosion [16,17]. Marsh aboveground vegetation builds elevation capital [18] by slowing water flow and facilitating surface sediment deposition, while belowground plant roots and rhizomes stabilize sediments, prevent erosion and contribute to building marsh elevation [19].

In the marsh interior, unvegetated patches can form, expand and coalesce leading to massive marsh dieback [20,21]. The genesis of these unvegetated patches can take multiple forms. Degraded marsh is less effective at accreting and building organic matter in the soil which can cause the marsh to lose elevation to a level outside of the growth range of marsh plants [19,22,23]. Such deterioration of the marsh facilitates further erosion and increased inundation triggering marsh dieback [19,24]. These poorly drained mud depressions, devoid of vegetation are called salt pannes. Salt pannes (hereafter ‘pannes’) are also referred to as salt pans [20,25], tidal flats, saline supratidal mudflats, salterns [26], pools [27,28], tidal ponds [29], or pond holes [10] (S1 Table). While many distinguish pannes from ponds and consider them to be two separate features of the marsh landscape [29,30], the latter retaining water and rarely draining and the former only inundated on the highest tides, in our study, we classify them collectively as ‘pannes’. Yap and colleagues first characterized the general morphology and dynamics of pannes in 1917 but noted that the factors that facilitate the original formation of pannes are poorly understood and likely vary across systems [20]. Pannes are thought to be formed by biogeomorphological processes [31] or physical stressors such as topographic depressions [32], tidal litter, waterlogging, or snow [33].

One potential driver of salt marsh dynamics at both bank edges and interior pannes is herbivory and bioturbation by crabs. Ubiquitous to most salt marsh systems, crabs have been shown to have strong yet variable effects on salt marsh structure and function. An observational study across fifteen US National Estuarine Research Reserves found that marsh cover is better predicted by elevation (and thus affected by sea-level rise) than crab or burrow abundance, though there are likely to be interacting effects between the two [34]. For example, in New England marshes, crab (*Sesarma reticulatum*) abundance has increased due to sea-level rise [35] and overfishing of predators [36]. This has led to runaway herbivory in the low marsh and changes to edaphic conditions induced by sea-level rise allowing crabs to move into previously inaccessible marsh [37]. In Argentina, the engineering of burrows by crabs (*Neohelice granulata*) facilitates the formation of salt pannes by lowering marsh elevation causing depressed patches to pool leading to marsh dieback and panne formation [31]. A study across three separate Southern California estuaries found that crab (*Pachygrapsus crassipes* and *Uca crenulata*) effects differed across sites and marsh plant species and when detected were positive [38]. Thus crab effects on marsh health likely vary temporally and spatially (across and within systems) and should be directly tested across a gradient of physical factors over time to gain a more complete understanding of potential effects to marsh health.

Most investigations of panne dynamics and of crab effects have occurred on the US East Coast, thus studies are needed elsewhere, both to seek generality across systems and to inform local management. We examined panne dynamics in Elkhorn Slough, an estuary located in Monterey Bay, California, where salt marsh loss has been documented at creek edges and the marsh interior [39], resulting in net loss of 70% of its historical salt marsh habitat [40], a major concern for regional stakeholders [41]. The majority of marsh loss in Elkhorn Slough occurs in the marsh interior through the formation and expansion of pannes [39]. Creek edge loss is affected by increased tidal velocities resulting from an artificial harbor mouth [42] and eutrophication [14]. Creek edges are riddled with crab burrows, constructed and maintained by native grapsid shore crab, *P. crassipes. P. crassipes* is an omnivorous crab that is found at its highest densities along creek edges and low marsh elevations [34]. Through both consumptive and engineering effects, *P. crassipes* has been shown to have strong negative effects on marsh plant biomass along creek edges, compromising the ability of the marsh to track sea-level rise and mitigate erosive forces (Beheshti et al. in review). Pannes were observed expanding during a period of severe drought (2012-2016) [43] and high water levels related to the warm water event known as “the Blob” (2013-2015) [44]. Panne dynamics remain poorly understood in the system and the majority of investigations [20,24,25,27–29] thus far have focused exclusively on physical factors with little attention paid to the role of crabs, although crab burrows are prominent along panne edges.

Our study investigates panne dynamics in Elkhorn Slough salt marshes and elucidates the role of crabs vs. other factors in driving marsh loss or recovery in the marsh interior. First, to explore the role of crabs on panne dynamics, we conducted a two-year field experiment where we attempted to manipulate crab densities across twenty pannes and evaluate panne response. We also explored how crab abundance and burrow density differed along creek versus panne edges using experimental data collected from studies conducted concurrently (2016-2018) at both areas of potential marsh loss. We failed to successfully manipulate crab densities and instead utilized the experimental plots as sub-samples to characterize the twenty pannes. We tracked the trajectory of pannes over the course of the study period by carefully monitoring the marsh-panne boundary where there is an abrupt transition from vegetated to unvegetated habitat. Elkhorn Slough experienced system-wide interior marsh loss in the years leading up to the study. Thus, the major emphasis of our investigation was to better understand which panne attributes (elevation, panne depth and size, distance to nearest panne, microphytobenthos, crab activity, and sediment dynamics) best predict panne expansion (marsh dieback) or contraction (marsh recovery). Our investigation will help inform the management of this estuary, as the first analysis of drivers of panne dynamics in a system that has experienced extensive interior marsh loss through panne expansion. Moreover, our study illustrates how integration of field data and modeling can elucidate the relative importance of multiple factors in driving marsh loss or gain, an approach applicable to any marsh system.

## METHODS

### Overview

We investigated dynamics at 20 pannes. Our study design involved experimental treatments manipulating crab densities replicated at each panne (S1 Fig). Since treatments were unsuccessful in affecting crab densities (see Supplemental Information), for later modeling of factors affecting panne dynamics, we used panne as replicate, averaging across treatment plots.

#### Study site

This study was conducted in Elkhorn Slough, an estuary located in Monterey Bay, California. Tides in the estuary are semidiurnal with a mean diurnal range of 1.7 m, a spring tidal range of 2.5 m and a neap tidal range of 0.9 m [45]. The mediterranean climate has temperatures averaging 11.1°C in the winter and 15.4°C in the summer [46]. The dominant marsh plant is *Salicornia pacifica* or pickleweed and the dominant grazer and bioturbator is *Pachygrapsus crassipes* or the lined shore crab.

#### Panne selection

We selected 20 pannes to study in a ~3 km stretch of salt marsh along the northwest side of Elkhorn Slough (Fig 1). Panne elevation was near Mean High Water (see details on elevations below) and the vegetation surrounding these pannes consisted almost entirely of the marsh dominant in this system, *S. pacifica*. Since our original focus was to conduct an experiment examining crab effects, we used criteria to select pannes that were physically similar to each other, to decrease variation in panne dynamics from other factors. We used geospatial analyses to select relatively similar pannes as follows (S2 Fig). Each panne had to 1) have been relatively stable in size from 2004 - 2012 as assessed in aerial imagery (chosen because we had access to high resolution imagery for these years), 2) be within 35 m from the nearest creek edge, 3) have a minimum 1 m buffer from the nearest panne, 4) have a diameter greater than 1.5 m, and 5) have either no secondary creek or if a secondary creek was present the width had to be less than 1 m. The chosen pannes that met the criteria described above had an average panne perimeter distance of 11.6 m (mean diameter = 3.7 m, standard deviation = 2.1 m) and the minimum and maximum panne perimeter distance was 8.16 m (diameter = 1.7 m) and 16.2 m (diameter = 5.1 m), respectively. Average distance to the nearest creek edge was 12.5 m (standard deviation = 8.5 m), the minimum and maximum distance was 2.13 and 34 m. Distance to nearest panne averaged 3.8 m (standard deviation = 2.3 m), the minimum and maximum distance was 1.18 and 11.4 m. The average secondary creek or ‘microchannel’ [47] width was 0.51 m (standard deviation = 0.28 m) and the maximum was 0.93 m, three of the pannes had no secondary creek. One of the pannes was discarded from the analyses because it was adjacent to a large seagrass meadow, resulting in extensive year-round wrack accumulation (resulting in final n=19).

**Fig 1.**
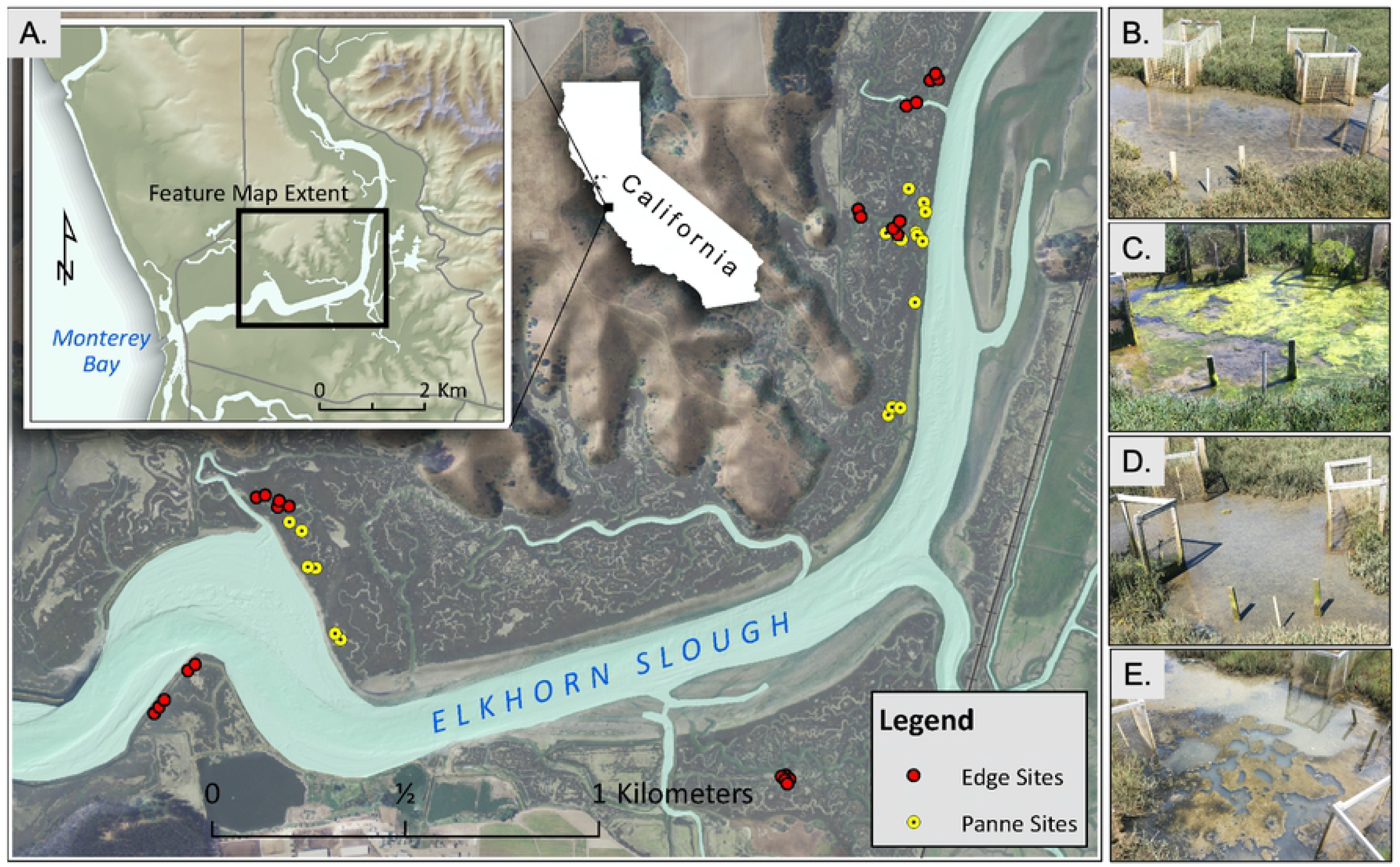
Study design from the landscape to plot scale. (A) Study map showing panne (yellow circles) and bank edge (red circles) studies. (B-E) Examples of the panne study and variation in panne size, pooling, and drainage.

#### Crab Experiment

Our original study design had one replicate of four treatments associated with each panne. The goal of the field experiment was to manipulate crab densities by means of fence design by either crab removals or additions (see Supplementary Information). These manipulations were ineffective at significantly altering burrow or crab numbers, and as a result we treated the 4 experimental plots in each panne as sub-samples of each panne (n=19). We derived variables measured at the plot level by taking the average across sub-samples per panne (e.g. burrow density was estimated as the average number of burrows of four sub-samples per panne). All other variables were taken at the panne scale (e.g. elevation, panne size and depth).

### Field data collection and indices

We collected data on movement of the vegetation edge, crabs, sediment dynamics and percent cover of succulent tissue and benthic algae within each of the four experimental plots in each of the 19 pannes.

#### Tracking panne expansion and contraction

To assess whether pannes expanded or contracted over the study period and at what rate, we installed five permanent transect line markers within each of the plots. Zip ties were used to mark the longitudinal start and end of each transect to ensure we were surveying the same points over time. During surveys, each of the transect lines was resurveyed and the last rooted vegetation along each transect line was recorded. Surveys were conducted annually from 2016 to 2018 (S3 Fig). To quantify contraction or expansion we calculated the average “marsh-panne boundary” difference per panne between 2016 and 2018. A positive value meant marsh colonization and panne contraction and a negative value meant marsh dieback and panne expansion. Our hypotheses for each of the parameters and indices outlined below can be found in S1 Table and S2 Table.

#### Crab Activity

To monitor crab density across pannes we conducted 24 hr crab trapping efforts annually (August 2016, March 2017, August 2018) within each of four plots per panne and visually assessed crab presence or absence. We monitored crab burrow densities annually to track any changes over time by counting all burrows over 1.0 cm in each plot (1.5 x 0.5 m), though burrows only occurred in the marsh zone of the plots (1.25 x 0.5 m). The ‘Crab Activity Index’ represents the mean number of burrows per plot (including both small, 1.0 cm - 2.9 cm, and large, 3.0 cm+ burrows) in 2018 and ‘Change in burrow density’ represents the relative change in burrow densities over the study period, both were included in the initial model. Previous work in Elkhorn Slough showed that the relationship between marsh biomass and crab engineering effects, measured as burrow density, are not the same as crab consumptive effects, measured as crab abundance (Beheshti et al. in review). Additionally, crab consumptive effects can be measured as either crab count, CPUE, or biomass. We included each of these measures of crab activity in the initial model (S2 Table).

#### Sediment Dynamics

To assess sediment dynamics in the panne and marsh of each experimental plot, we installed galvanized conduit rods (3.048 m-long with a 1.905 cm diameter) in the panne and marsh zone of each of the four plots per panne. Rods were installed using a ladder and post driver until we reached hard ground, this typically occurred around 2.75 m, leaving around 30 cm of rod exposed (Video S1). More of the rod exposed over time indicated erosion or compaction and less of the rod exposed over time, accretion or expansion. The change in rod exposed was calculated using the following equation: Δ_rod_= (Rod_2016_ - Rod_2018_). The ‘Panne Sediment Dynamics Index’ and ‘Marsh Sediment Dynamics Index’ represent the mean change (from 2016 to 2018) in panne and marsh rod exposed, respectively. Both were included in the initial model (S2 Table).

#### Percent cover of succulent tissue and benthic algae

To determine the potential role of aboveground productivity on panne dynamics, we evaluated the change in new succulent growth over the study period (2016-2018), hereafter termed the ‘Marsh Productivity Index’. In pickleweed marshes, succulent tissue is new growth that represents the present growing season and the woody tissue is older growth from years prior [48]. We were interested in the relative changes to succulent growth as a proxy for marsh productivity; a relative increase in succulent cover would indicate potential marsh recovery (S2 Table). To assess percent cover of succulent tissue, we placed a 50 x 50 cm gridded quadrat in the middle of the marsh portion of each plot and dropped a metal rod at 20 intercepts. The number of succulent cover intercepts was divided by the total possible points (n=20) and multiplied by 100 to get succulent tissue percent cover. This was done in 2016, 2017 and 2018 at each of the four replicate plots per panne. Here we present data comparing the absolute differences for each response variable between 2016 and 2018 only.

To determine the potential role of benthic algae, a catch-all category that included diatom biofilms and macroalgae (*Ulva* sp., *Vaucheria* sp., etc.), on panne dynamics, we evaluated the relative change in benthic algae over the study period (2016-2018). We assessed benthic algae cover, hereafter termed the ‘Biofilm Index’, using the same methods described above. We were interested in the relative changes (2016-2018) to the Biofilm Index and the possible relationship between the Biofilm Index and panne contraction or expansion (S2 Table). This was done in 2016, 2017 and 2018 at each of the four replicate plots per panne, we are presenting the absolute differences between 2016 and 2018 only.

### Geomorphological data collection and indices

We used geospatial analyses to characterize potential factors that might affect marsh dynamics at each of the 19 pannes.

#### Indices for panne depth, elevation, size, and distance to nearest panne

Using a LiDAR Digital Elevation Model (2018), we estimated the elevations for all nineteen pannes that were used in analyses. To calculate panne depth, we used ArcGIS v. 10.7 and 2018 NAIP 4-band orthoimagery (upgraded to 15 cm resolution) to create polygons of each panne (S4 Fig). We applied a 1m buffer to the polygons and used the panne/buffer mask to extract cell values from the 2018 LIDAR (1 m resolution) using the Spatial Analyst Zonal Statistics as Table tool. Depending on size, between 12 to 37 cells per panne were used to compute the minimum, maximum, range, and mean elevation values (NAVD88, meters) of each panne (n=19). LIDAR elevations were corroborated by real-time kinematic positioning (RTK) at five experimental plots. We did not detect a significant positive elevation bias due to vegetation [49]. Panne depth was measured as the elevation difference between a single elevation point in the middle of the panne (panne elevation) and the surrounding marsh (non-vegetated vertical accuracy between 8-10 cm).

Pannes are typically circular in shape but can be irregular. The ‘Panne Size Index’ represents the panne perimeter distance. The perimeter of each panne (at the last rooted vegetation) was carefully traced using a transect tape. Using the same transect tape we measured the shortest possible distance to the nearest panne (‘Distance to Nearest Panne’), the width of any branching microchannel in the panne (‘Microchannel width’), and the shortest distance to the nearest tidal creek (‘Distance to bank edge’) (S2 Table).

### Comparison of crab and burrow densities along panne and tidal creek bank edges

To compare crab and burrow densities in panne vs. creek edges, we summarized data at the block level for panne (n=19) and creek (n=25) edges (Beheshti et al. in review). Crab abundance data from both experiments was compared by calculating the average crab Catch Per Unit Effort (CPUE), with effort being a single sampling unit or pit-fall trap. Crab CPUE was averaged across 8 pit-fall traps for the panne study and 4 pit-fall traps for the creek study. Burrow densities were averaged across 4 replicate plots for the panne study, across 2 replicate plots for the creek study. Burrow densities for both panne and creek plots were expressed as density per m^2^. Data used to compare crab effects across both studies were collected in August 2018.

### Statistical analyses

We examined the role of a suite of physical and biological factors (S2 Table) in explaining the rate of marsh recovery or dieback along panne edges. To do this, we summarized the data at the level of panne (n=19), using the four plots in each block as sub-samples. We then used a stepwise regression to determine which model effects were predictive of panne contraction or expansion. Final model selection was based on Akaike Information Criterion (AIC). To compare crab CPUE and burrow densities between creeks and pannes, we used an ANOVA (two levels of factor “Location”; Panne and Creek Edges).

## RESULTS

### Characterization of panne trajectory and indices

Overall we found that pannes contracted during the study period. The movement of the vegetation boundary towards the panne center averaged 16.30 ± 7.83 cm/year (minimum = 6.22 cm, maximum = 33.79 cm). Recovery appeared to consist entirely of clonal expansion by existing *S. pacifica* growing around the panne edge; we did not observe plants colonizing the panne area via seed.

### Model results identifying correlates of panne recovery rate

In our initial multiple regression model we checked the Variance Inflation Factor (VIF) and observed high (>10) VIF scores for elevation and depth, which was inappropriate for the model. After looking at the covariance structure between panne depth and elevation, we found that the two were highly inversely correlated (S4 Fig; lower elevation associated with deeper pannes and higher elevation associated with shallower pannes). Therefore, we developed a Principle Component variable combining these two variables, hereafter PC1 (Depth and Elevation) to include in the stepwise regression. Higher values of PC1 represent higher elevation and shallower pannes and lower values of PC1 represent lower elevation and deeper pannes. None of the other variables showed significant covariance; they each had low VIF scores (<2).

The stepwise regression identified the best predictive model out of the initial 14 parameters included (S1 Table). The final model included 5 model effects: Panne Size Index, PC1 (Depth and Elevation), Distance to Nearest Panne, Sediment Dynamics Index, and Crab Activity Index (Table 2). All model effects in the final model were significant and had VIF scores less than 2.0. For definitions of the aforementioned indices see Methods and Table 1.

**Table 1.**
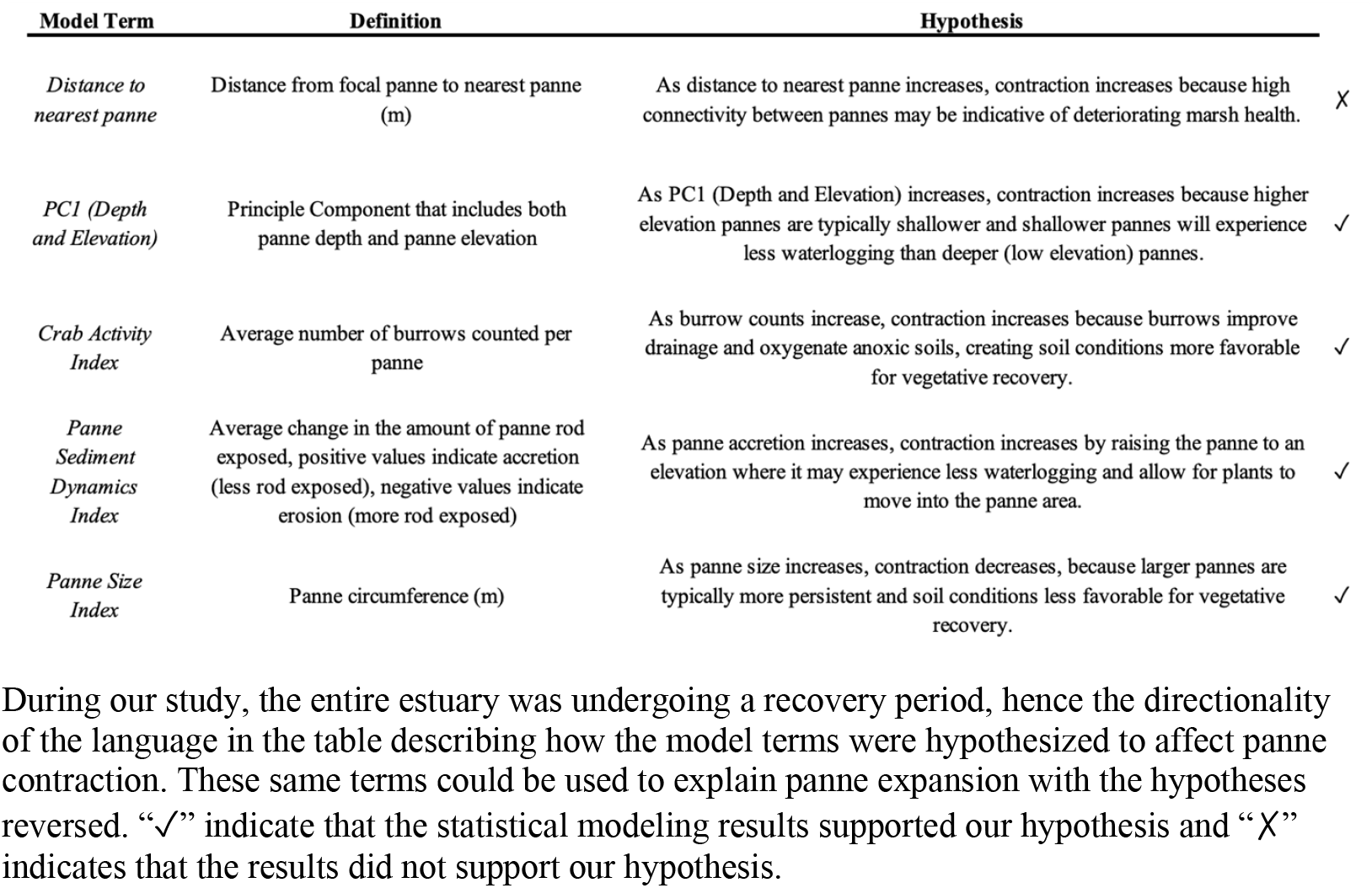
Model terms, definitions and hypotheses about panne contraction.

The overall fit of the model was high, with ~86% of the variation in the response (absolute movement of marsh-panne boundary (cm) from 2016-2018) explained by the model (R^2^=0.857, F_5,13_=15.664, p<0.0001). Panne size was negatively associated with panne contraction (Slope = −1.169), the larger the panne, the lower observed panne contraction. PC1 (Depth and Elevation) was positively associated with panne contraction (Slope = 4.559), with higher elevation and shallower pannes contracting more than low elevation deeper pannes. Nearness between focal pannes and adjacent pannes was negatively associated with panne contraction (Slope= −3.289), with larger distances between pannes correlated to less contraction. Crab burrowing activity was positively correlated with panne contraction (Slope= 0.315), as burrow density increased, so did panne contraction. Lastly, sediment dynamics was positively associated with panne contraction (Slope=1.788); pannes that accreted contracted more than pannes that showed no change over the study period or eroded (Table 2, Fig 2).

**Fig 2.**
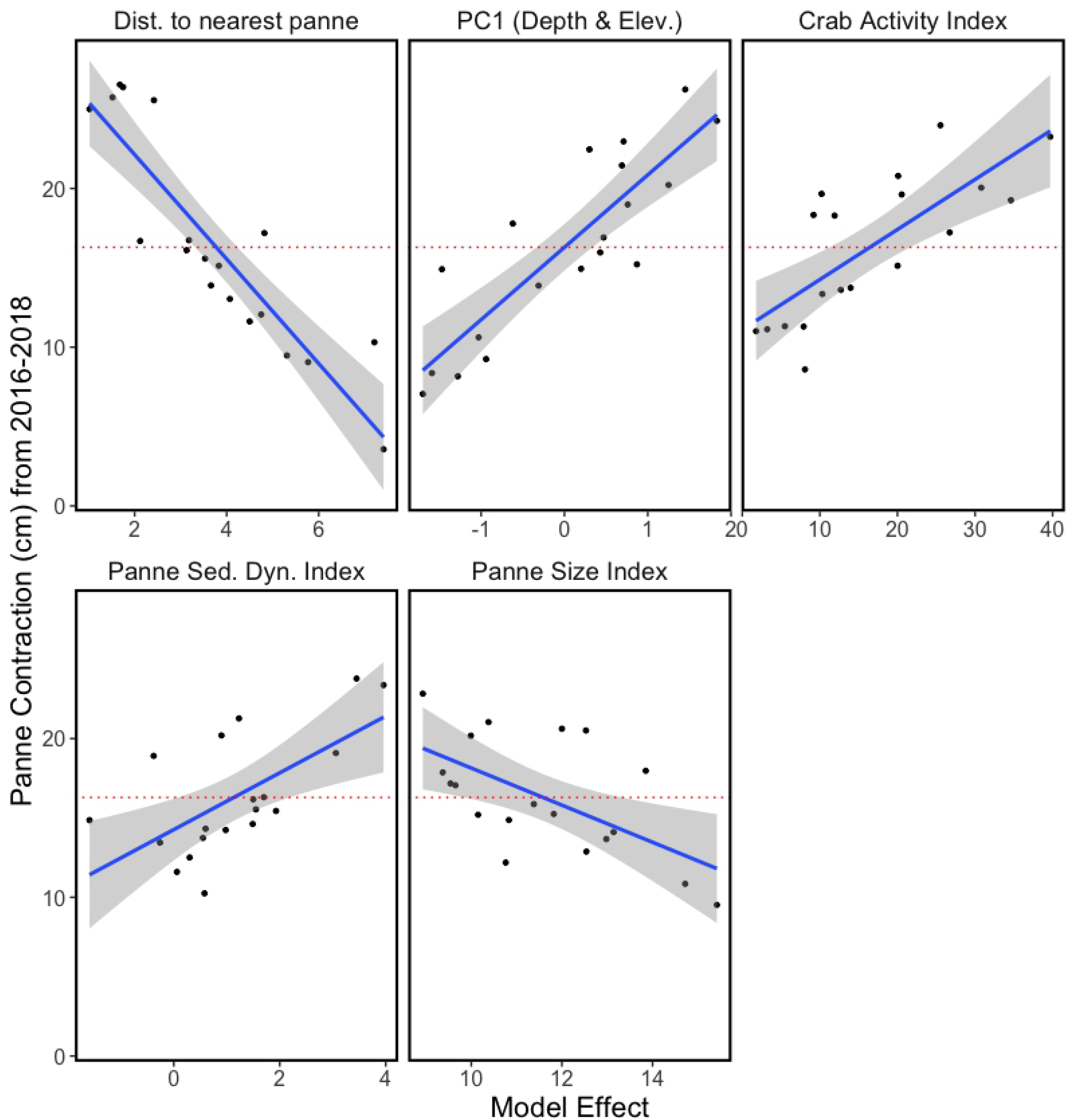
Partial leverage plots for all of the best-fit model effects. Plotted is the movement of marsh into the panne area (panne contraction along marsh-panne boundary) in cm. The dotted red horizontal line represents the average marsh-panne boundary movement from 2016-2018 of 16.298 cm. Each partial leverage plot includes the 95% C.I. For a list of the parameters included in the plotted indices, see Table 1.

**Table 2.**
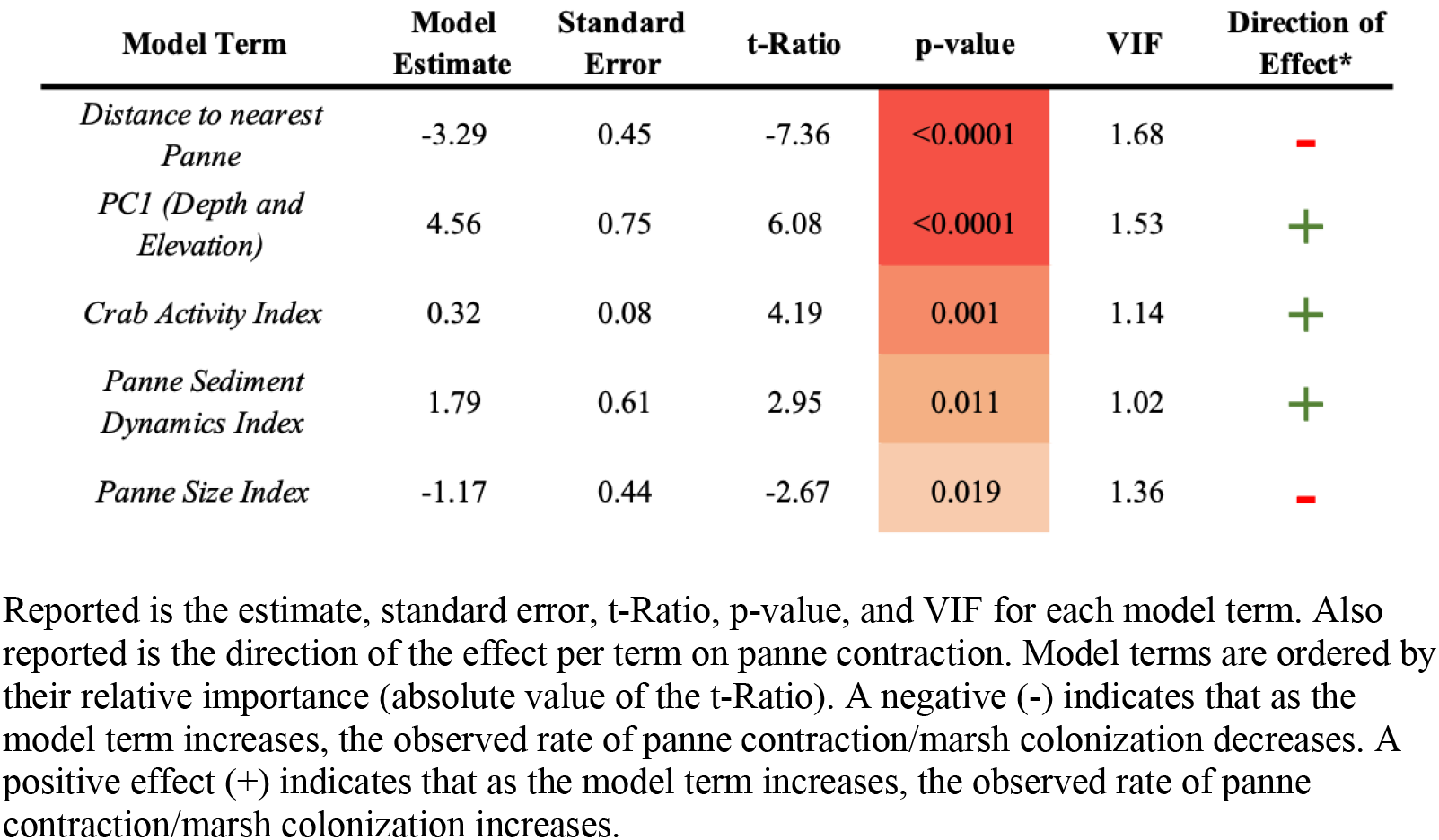
Multiple regression model output.

### Comparison between crabs at bank edge vs. pannes

Crab CPUE was significantly greater along creek edges relative to panne edges (ANOVA; F_1,42_=30.53, p<0.0001) (Fig 3A). Additionally, burrow density was significantly greater along creek edges relative to panne edges, with approximately 4x as many burrows observed along creek edges (ANOVA; F_1,42_=25.15, p<0.0001) (Fig 3B).

**Fig 3.**
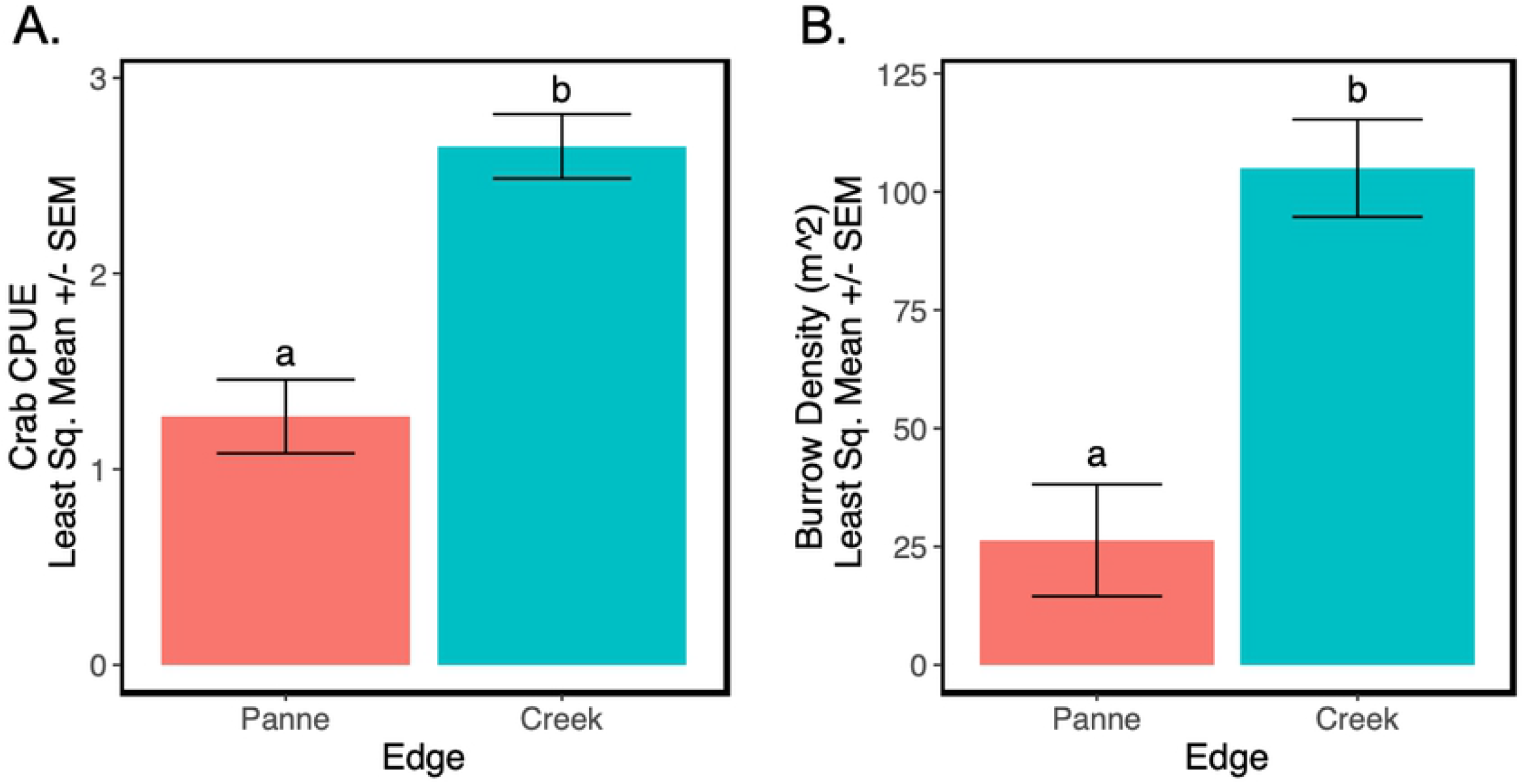
Comparison of crab CPUE and burrow density along panne versus creek edges. (A) Crab CPUE along panne versus creek edges. (B) Burrow density (# per m^2^) along panne versus creek edges. Plotted is the Least Square Means Estimates ± Standard Error. Pink bars represent panne edges and blue bars represent creek edges. Different letters denote significant differences (□ = 0.05) between panne and creek edges.

## DISCUSSION

### Multiple local factors drive interior marsh dynamics

Recently, Zhu and colleagues called pannes the “unrecognized Achilles’ heel of marsh resilience to sea-level rise” [50]. Much marsh degradation results from panne formation and expansion, but the mechanisms behind panne dynamics are not broadly understood. Some seminal studies have characterized key drivers [10,27,33,47,51]. Our investigation complements this earlier work and provides the first study of panne dynamics in California marshes, which are dominated by a perennial succulent, different from many of the other study systems (i.e. herbaceous grass-dominated marshes). Additionally, while there have been multiple studies demonstrating that salt marshes are structured by both physical and biological factors and their interactions [35,52,53], most investigations of panne dynamics have focused almost entirely on geomorphology [32,46,59; for exception see 30]. In exploring how physical (panne attributes and sediment dynamics) and biological (crab activity) factors affect marsh recovery along panne edges, our study provides a novel perspective on drivers of panne dynamics along the US west coast.

Salt panne dynamics are strongly controlled by drainage [10,20,33]. In Plum Island Estuary (Massachusetts, USA), salt marsh has kept pace with sea-level rise while panne area has increased and drainage density decreased, suggesting that drainage is a stronger driver of panne dynamics than sea-level rise [47]. Over a two-year period, an experimentally drained panne did not change in depth but marginal revegetation did occur on exposed mud with the alleviation of waterlogging stress (i.e. anoxia, sulfide toxicity, hypersalinity) [47]. In our study we saw similar patterns. In the most supported model, PC1 (Depth and Elevation) was positively associated with panne contraction, with marsh recovering relatively quicker at shallower high elevation pannes than deeper low elevation pannes (Fig 4, S5 Fig). Deeper pannes pool and retain water after tidal inundation more than shallow pannes. Previous work has shown that panne formation may be driven by depth and inundation time [30]. Elkhorn Slough was experiencing system-wide marsh recovery during our study period, and in the marsh interior, our study suggests that the potential for marsh recovery is greatest at shallower, high elevation pannes where drainage is likely higher. This may due to similar mechanisms as those observed in Plum Island Estuary, where waterlogging stress observed in deeper pannes inhibited marsh recovery into the panne area.

**Fig 4.**
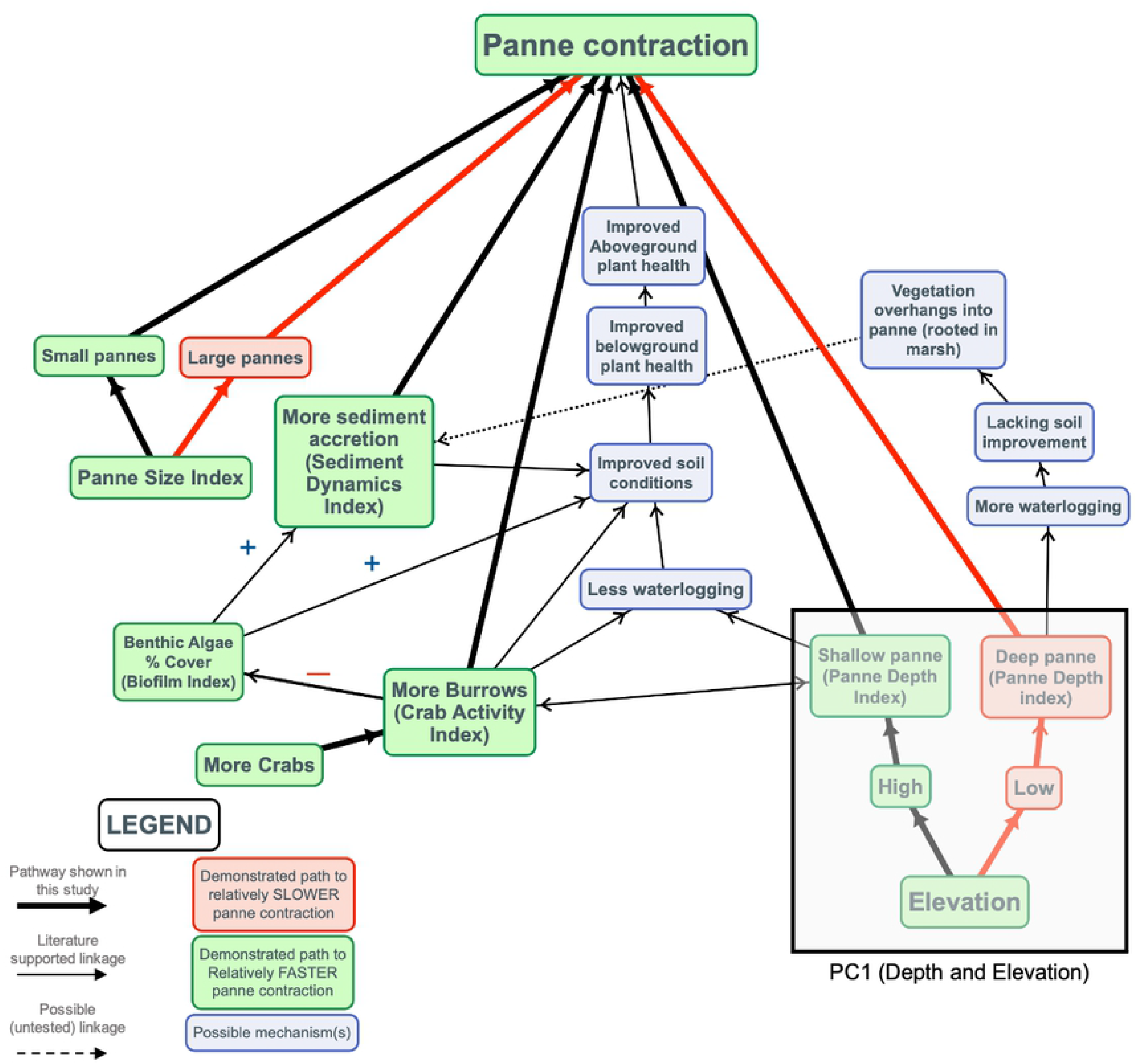
Conceptual model of panne contraction and possible mechanisms. Bolded arrows indicate relationships or main model terms that correlated to panne contraction and were shown in our study. Narrow arrows indicate pathways that were not demonstrated by our study but are likely to play a role based on marsh dynamics literature. Red arrows and cells indicate pathways to slower panne contraction and green cells indicate pathways to relatively rapid panne contraction. Blue cells indicate potential mechanisms.

Panne size has also been shown to be critically important in predicting the trajectory of marsh recovery. Smaller pannes are prone to infilling resulting in marsh recovery while larger pannes experience wave-induced erosion and bed shear stress which causes pannes to deepen and expand [52] often coalescing into other pannes [20]. Our results complement previous work by showing greater rates of panne contraction for smaller versus larger pannes (Fig 4).

Panne density has been shown to be negatively correlated to creek density [33], further supporting the hypothesis that poor drainage (fewer creeks) promotes panne formation and persistence. Our results differ from those of Goudie [33] and indicated that marsh recovery was greatest when pannes were near to one another (Fig 4). While we expected regions with high densities of pannes to signal poor marsh health, our data suggests that recovery is more rapid in regions where pannes are closer to one another suggesting that there may be inter-panne effects at play (i.e. sediment exchange, improved drainage from panne to panne, etc.).

Sediment dynamics in the pannes themselves is also of great importance and affected by panne elevation [19,24], depth [47], size [28], and crab burrows (this study). The results of our study also showed that local increases in marsh elevation (due to panne accretion or soil expansion) was positively correlated with panne contraction. Conversely, the pannes with local marsh elevation loss at the marker rods showed the lowest rates of marsh recovery. Accretion can improve soil conditions by possibly both raising the panne to an elevation within the growth range of marsh plants and alleviating stressors associated with waterlogging (Fig 4). Our results mirror previous work by showing an association between accretion and marsh recovery [47]. We suspect that increased elevation and improved drainage are both necessary for marsh colonization into pannes, as it is likely that an erosional panne that is well drained will only deepen and ultimately reach a tidal elevation below the growth range of marsh plants [19,33,47,53].

While crab burrows have been shown to have positive effects on marsh productivity, mainly by oxygenating anoxic sediments [54] or increasing nutrient uptake [55], crab burrows have also been found to increase erosion and creek formation [56,57] and elongation [58]. In Argentina marshes, crab burrows facilitate the formation of pannes through loss of elevation (see Fig. 9 in 30). In our study, we observed crab burrows having a positive effect on panne contraction (Fig 4). In Elkhorn Slough, it is likely that the positive effect of burrows is due to increased drainage [58] and reduced waterlogging stress [59], which outweighs any possible negative effects.

### Crab effects are context-dependent

Based on the results of this and previous studies, effects of *P. crassipes* on marsh dynamics in Elkhorn Slough are context-dependent, with different physical factors across the marsh landscape changing not only the strength but the direction of certain crab effects. Crab burrows were found to be negatively associated with marsh biomass along creek edges (Beheshti et al. in review). In the current study we found that burrows are positively associated with marsh recovery and panne contraction. The positive association between burrows and panne contraction is likely linked to improved drainage and an indirect effect on soil improvement (i.e. oxygenation of anoxic sediments, less sulfide buildup) (Fig 4). The different direction of crab effects and spatial differences in abundance are likely due to different physical factors driving dynamics along creek versus panne edges. For example, creek dynamics are driven by erosive processes [2] and have different hydrodynamics and geomorphology compared to panne edges. This contrast highlights how complex and variable the geomorphology can be in salt marsh systems where pannes in close proximity (~2 to 34 meters in our study) to creek edges can have entirely different relationships between the same physical and biological drivers.

Top-down effects on vegetation can be very strong [61–63], but such effects will always interact with physical factors and thus may vary in strength (i.e. snail grazing in marshes with drought [64], pollinator and herbivore interactions with plants across environmental gradients [65], foraging behavior of coral reef fishes with distance from reef [66]). Crab effects on salt marshes have been shown to have such context-dependent variation when examined across different estuaries [34,38,67]. In a meta-analysis that included up to 42 studies assessing consumer effects of crabs on salt marsh plants, the average effect size (Hedges’ g) for multiple response variables (e.g. above and belowground biomass, plant survival, density) was overwhelmingly negative (n=50) as opposed to positive (n=8) (see Fig. 4 in He and Silliman 2016) [67], demonstrating that crab effects, though typically negative, are not uniform and are instead context-dependent. In New England marshes, burrowing by crab *Uca pugnax* were shown to increase drainage and redox potential in the sediments of cordgrass marshes, promoting biomass production in soft sediment marsh environments [54]. In another example showing positive effects of crabs on marsh vegetation, Holdredge and colleagues [55] found that in sandy cordgrass marshes, crabs positively affected nutrient uptake by cordgrass. They also found that experimentally removing crabs caused above- and belowground biomass to drop by ~50% [55]. These two studies show that the mechanisms driving positive crab effects differ for marshes with different physical characteristics (e.g. fine vs coarse sediment). In other studies, in Argentina marshes, the effects of burrowing by crab *Neohelice granulata* varied across the marsh landscape, promoting sediment trapping in the marsh interior (positive effect) and enhancing sediment transport on creek edges (negative effect) [56] and in a separate study, consumer pressure by crabs was shown to prevent marsh colonization of pannes [63].

### Regional and global drivers of marsh dynamics

While local factors and attributes of the pannes themselves predict short-term panne dynamics, it is clear that regional and global drivers also can exert strong effects. Our study was conducted immediately following one of the worst droughts in California history [43] and a warm, high water event [44] that coincided with a period of marsh loss and panne expansion in Elkhorn Slough. Following these dry and warm periods was the second wettest season (2016-2017) in California since 1951 [68]. Precipitation may play an important role in facilitating the recovery of the marsh along these physically stressful panne edges in mediterranean marshes prone to high salinities. Further investigation is needed to understand how regional climatic and oceanographic events affect panne dynamics.

Our study showed how elevation, a proxy for relative sea-level rise, can affect rates of panne contraction. We tracked relatively slower panne contraction at low elevations, suggesting that opportunities for marsh recovery are diminished with sea-level rise. We also found that local increases in elevation at our marker rods correlated with panne recovery. Some studies have indicated that sea-level rise will increase the rate of panne formation, expansion, and coalescence, further contributing to marsh loss [24,50], while others suggest that poor drainage, insufficient accretion, and poor creek connectivity explain panne formation [47]. In our study we show that elevation, drainage, and inundation are inextricably linked and predictive of panne dynamics. Further, as sea-level rise continues, channels may deepen and widen, increasing the tidal prism [33] and crabs may become more abundant as low marsh extent increases [35]. This has already been demonstrated in New England marshes [35,58]. The interaction between crab effects and sea-level rise on panne dynamics needs further study. Our results indicate that crab burrowing along panne edges facilitates marsh recovery and panne contraction, likely by improving drainage and reducing waterlogging. As a reminder, our study was initially designed as a crab experiment and we specifically chose pannes that were relatively stable and did not vary greatly in geomorphology to reduce the possible confounding factors that may have impacted our ability to detect treatment effects. This makes our findings all that more compelling considering that we were able to identify predictors of panne contraction across 19 pannes with slight physical differences. We recommend future studies encompass the actual variation in panne characteristics seen across the estuary and expect patterns shown here to only strengthen with the inclusion of pannes of a wider range of sizes, shapes, elevations, and statuses. Relatedly, further study is needed to explore how the factors identified in this study may change with projected sea-level rise, one way to approach this would be to include pannes from a wider range of marsh elevations, as a better proxy for sea-level rise (S6 Fig).

Our understanding of panne dynamics is improving, but more robust predictions are needed of how dynamics may shift with both short and long term disturbances associated with global change. Future work should track panne formation, expansion, coalescence, or contraction as sea-levels rise and anthropogenic stressors worsen [9,30] and compare rates of interior marsh loss to historical rates to improve the management of these coastal habitats and inform marsh conservation strategies. Such studies are needed to better understand how extreme meteorological and oceanographic events, such as those that preceded and occurred during our study, affect marsh dynamics and resilience.

## ACKNOWLEDGEMENTS

We would like to thank UCSC undergraduate interns A. Lapides, H. Heigl, C. Oliveira De Alcantara, J. Ruud, S. Kawamoto, L. Hijikata, N. Dwyer, and A. Pacheco and high school interns H. Levy, M. Levy, and J. Schwartz for their field assistance. We thank P. Raimondi for his guidance and assistance with data analysis and for sharing his enthusiasm for the principles of ecology and the stats that tell the data story, and J. Haskins and A. Woolfolk for sharing their expertise on the hydrology of pannes and the use of pannes by crabs. Lastly, we would like to thank R. Eby for his assistance with the design and installation of the fences initially intended for this study and for boat support.

## Supporting Information

**S1 Table. Examples of the different terminology used to describe pannes in the literature.**

All of the terms listed in the table are collectively referred to as ‘pannes’ in our study.

**S2 Table. Full list of model terms and direction of significant effects.**

Non-significant terms were excluded from the final model and are denoted with “ns”. The directionality of effect for non-significant terms is the hypothesized directionality, the directionality of effect for significant terms is model-based. Some hypotheses (e.g. ‘Distance to Nearest Panne’) were the opposite of what we observed. *A negative effect (-) indicates that as the model term increases, the observed rate of panne contraction/marsh colonization decreases. A positive effect (+) indicates that as the model term increases, the observed rate of panne contraction/marsh colonization increases.

**S1 Fig. Photos of study design.**

(A) Study design with treatments labeled at single block, (B) Above Ambient Crab experimental plot with the flashing installed flush to the fence wall, (C) *P. crassipes* crab in a burrow in one of our experimental plots and (D) Close up view of the panne rod and the zip-tie marker for the transects that run from the panne-edge to marsh-edge of the plot (See S3 Fig).

**S2 Fig. Heat map of salt marsh zones.**

Areas where there has been net gain (blue) or loss (red) of habitat from 2004-2012. Areas of high gain or loss (dark colors) were not included in this study.

**S3 Fig. Schematic of how marsh-panne boundary was monitored over time in fenced plots.**

Changes to the marsh-panne boundary over time indicate either marsh colonization and panne contraction (as pictured here) or marsh dieback and panne expansion.

**S1 Video. Installation of galvanized conduit rods.**

Rods were installed into the panne and marsh area of each experimental plot across all twenty blocks, or pannes. Link: https://youtu.be/nHdfJg4o-Bw

**S4 Fig. Example of panne polygons used to determine panne depth.**

A 1 m buffer circle (shown in yellow) was used to extract the 2018 DEM cells. Raw lidar points (shown in the image as green points) were not used since they were not particularly well-spaced. The DEM uses an interpolation between the points, and thus was reliable at representing the marsh-panne boundary and the pannes themselves.

**S5 Fig. Relationship between elevation and panne depth.**

This inverse correlation led to the development of a Principle Component (PC1 (Depth and Elevation)). Deep pannes are indicated by black circles and shallow pannes by blue triangles. Reported in the top left corner of the plot is the R^2^ and plotted regressions include the 95% C.I.

**S6 Fig. Elevation by panne size.**

As panne elevation increases, size decreases (R^2^=0.15). Similar patterns were observed in Escapa et al. [31] (See Fig. 3)--panne size (‘Patch diameter’; Escapa et al. 2015) decreased as elevation increased. The lower elevation edge for pickleweed in Elkhorn Slough is ~1.20 m NAVD 88 (*C. Endris, unpublished data*), our study did not extend lower than 1.37 m NAVD 88.

**S7 Fig. Experimental results.**

(A) crab CPUE and (B) change in marsh-panne boundary (2016-2018) by treatment and elevation. High elevation blocks are plotted in orange and low elevation blocks in blue.

